# EquiFold: Protein Structure Prediction with a Novel Coarse-Grained Structure Representation

**DOI:** 10.1101/2022.10.07.511322

**Authors:** Jae Hyeon Lee, Payman Yadollahpour, Andrew Watkins, Nathan C. Frey, Andrew Leaver-Fay, Stephen Ra, Kyunghyun Cho, Vladimir Gligorijević, Aviv Regev, Richard Bonneau

**Affiliations:** Prescient Design, Genentech; Genentech Research and Early Development, Genentech; Department of Computer Science, Courant Institute of Mathematical Sciences, New York University; Center for Data Science, New York University

## Abstract

Designing proteins to achieve specific functions often requires *in silico* modeling of their properties at high throughput scale and can significantly benefit from fast and accurate protein structure prediction. We introduce EquiFold, a new end-to-end differentiable, SE(3)-equivariant, all-atom protein structure prediction model. EquiFold uses a novel coarse-grained representation of protein structures that does not require multiple sequence alignments or protein language model embeddings, inputs that are commonly used in other state-of-the-art structure prediction models. Our method relies on geometrical structure representation and is substantially smaller than prior state-of-the-art models. In preliminary studies, EquiFold achieved comparable accuracy to AlphaFold but was orders of magnitude faster. The combination of high speed and accuracy make EquiFold suitable for a number of downstream tasks, including protein property prediction and design.

## 1 Introduction

Recent studies using deep neural networks to predict protein structure [1, 2, 3, 4, 5] have accelerated the development of structure-based methods for protein property prediction and design. However, some models tend to predict one or few conformations that may not be optimal for properties of interest, such as ligand-binding pockets [6] and protein-protein interfaces [7, 8, 9]. Moreover, these models require either multiple sequence alignments (MSA), protein language model embeddings, or other statistical information distilled from large sequence databases. While complex inputs such as embeddings and MSAs have proven useful for structure prediction, they increase the complexity of properly testing these methods, require significant time to derive, and often scale poorly with respect to sequence library size, and thus may overall limit downstream use. Conversely, the physics of protein dynamics and interactions do not depend directly on these inputs.

Here, we introduce EquiFold, a novel representation of protein structures and an end-to-end differentiable, SE(3)-equivariant neural network that predicts a protein structure given its primary sequence via iterative refinement. EquiFold makes atomically accurate predictions for *de novo* designed miniproteins with in *silico* predicted structures [10] and on experimental antibody structures from the Protein Data Bank [11]. We focus on this set because small structures (e.g., designed mini-proteins) and flexible loops (e.g., the CDR-H3 loop of IgG based antibodies) have proven to be as difficult to predict as larger structures in recent tests of state-of-the-art methods [12, 13, 14]. The model relies solely on geometrical structure representation and can readily incorporate various energy functions as physical priors, which we hypothesize to be instrumental in exploring conformational landscapes towards predicting properties of interest.

## 2 Related Work

AlphaFold [1] and similar models [2, 3, 4] have been successful at protein structure prediction, by representing the geometry of the chain of backbone atoms as a set of nodes paired with Euclidean transformations and using an iterative refinement procedure that updates the transformations per each block of the structure module. In these models, side-chain geometries are implicitly modeled until they are predicted as a series of torsion angles by the module’s final block. However, such implicit modeling of side-chains may make it difficult to model their atoms’ placements and interactions in 3D space. For example, to avoid steric clashes, such models must learn from data complex distributions in the high dimensional space of torsion angles that span multiple residues. In contrast, models that represent side-chain degrees of freedom explicitly in 3D would likely need to learn substantially simpler distributions to achieve the same goal.

Coarse-grained structure representations of proteins [15, 16, 17] typically model each residue by one or few nodes with associated positions determined by its atoms’ coordinates. These strategies can increase computational efficiency for predictive tasks, such as interacting residue predictions [18] and functional residue prediction [19], and are appropriate for certain generative modeling tasks, such as generating backbone scaffolds [20, 21]. However, these approaches have previously sacrificed all-atom structure resolution needed for design and packing-related tasks, along with much information useful for predicting protein functions.

To address these limitations, we develop a novel coarse-grained representation that retains all-atom structure resolution. In this representation, side-chain degrees of freedom are modeled explicitly in 3D space rather than intrinsically through torsion angles, which we conjecture makes it easier to model geometry and interactions in 3D space.

SE(3)-equivariant neural networks have been used in various 3D object modeling tasks, including atomic potential prediction [22, 23, 24], molecular property prediction [25, 26, 27, 28, 29], protein structure prediction, [2] and docking [30, 31]. They incorporate the symmetry of 3D space and are substantially more data efficient than their non-symmetry-aware counterparts [32, 33]. Input to SE(3)-equivariant networks consists of geometric tensors, or the irreducible representations of the SO(3) group (the group of rotations in 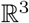), of various degrees *l*, such as scalars (*l* = 0), vectors (*l* = 1), and higher degree (*l* ≥ 2) tensors that transform under a rotation *R* through multiplication with corresponding Wigner D-matrices *D_l_*(*R*). Objects with an associated rotation, such as backbone frames [1], can be initially embedded with geometric tensors [15]. We use an SE(3)-equivariant model adapted from Equiformer [29] along with an initial embedding of coarse-grained nodes with geometric tensors.

## 3 Methods

### 3.1 A coarse-grained representation

Given a protein sequence *a* = (*a*_1_,…, *a_N_*) of length *N* where *a_i_* is one of the 20 canonical amino acids, its structure is specified by the set of 3D coordinates of its atoms grouped by amino acid, 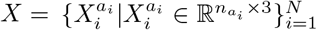 where *n_a_i__* is the number of atoms in residue *a_i_*. In a CG representation, each amino acid *a_i_* is represented by a predetermined set of CG *nodes* 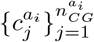, where each CG node 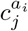 represents a subset 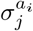 of the amino acid’s constituent atoms. The CG nodes are chosen such that 1) their union represents all atoms of the amino acid; 2) each member atom of a node shares at least one covalent bond with another member of the same node; and 3) each node consists of at least three atoms that collectively form a rigid body whose orientation in 3D is uniquely determined. Based on the last property, we define a *forward* CG mapping 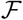 of the 3D coordinates 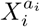 of an amino acid *a_i_*’s atoms into its CG representation:

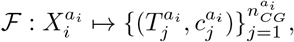

where each tuple in the set consists of a CG node identity 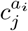 and a corresponding Euclidean transformation 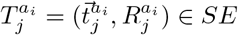(3) that maps the predefined template coordinates for atoms in 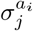 to the corresponding input atom coordinates (see Appendix A.1). Table 3 illustrates an example of a CG scheme used in this work that respects the above properties while minimizing the redundancy of atom representations across CG nodes. Figure 1 gives illustrations of this CG scheme for select amino acids.

**Figure 1:**
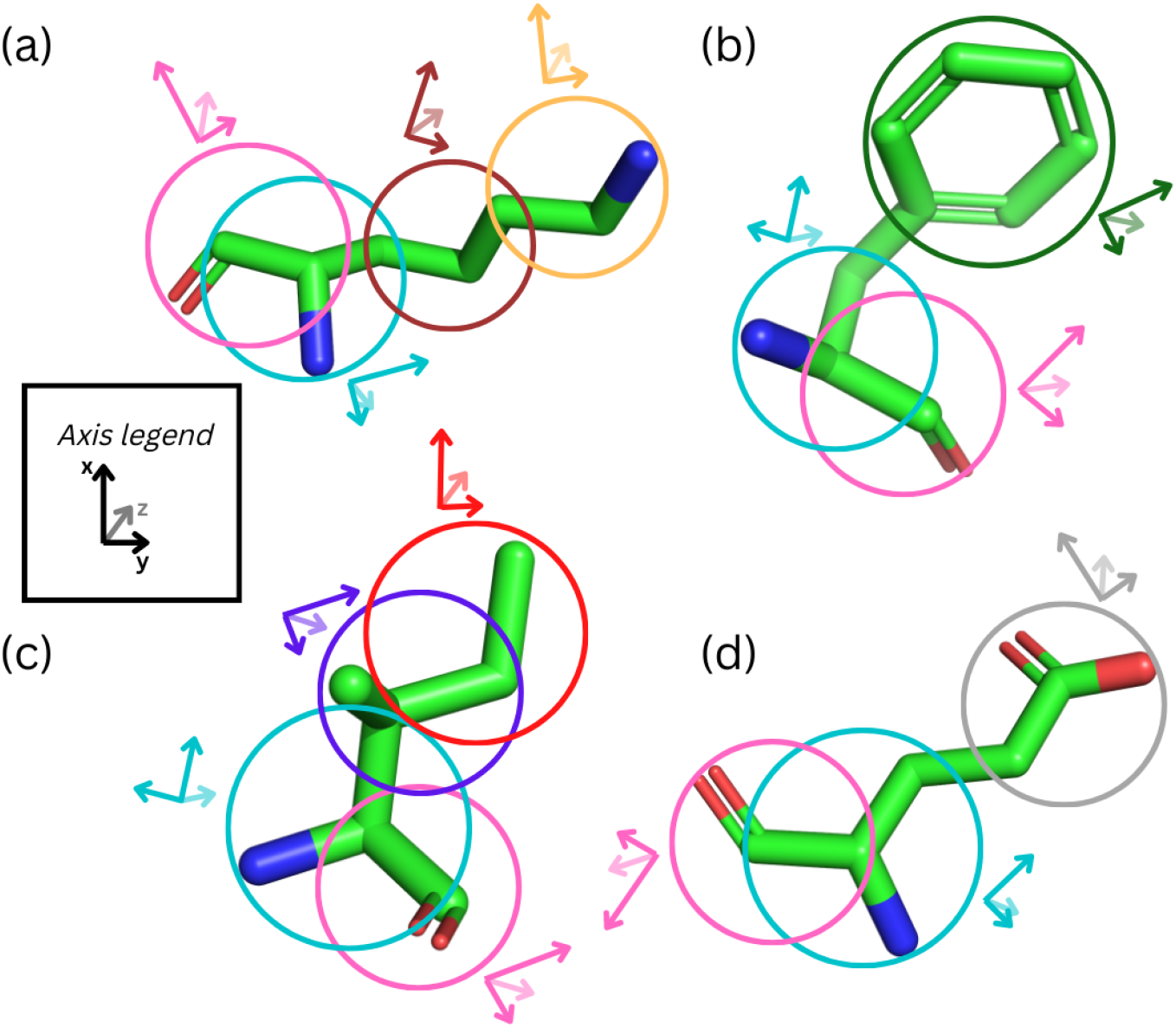
A representation of EquiFold’s coarse-grained node scheme as applied to four amino acids: a) lysine, b) phenylalanine, c) isoleucine, and d) glutamate. Atoms belonging to each node are circled and axes representing the node’s specification (by rotation-transformation) are shown in a corresponding color. Backbone nodes, which are common to each amino acid, are shown in consistent colors across panels.

**Table 1:** Performance of EquiFold on *de novo* designed mini-proteins from the test set broken down by fold. All RMSDs are reported in angstroms. “RMSD” and “*C_α_* RMSD” report all atom and backbone *C_α_* atom RMSDs based on the comparison of EquiFold predictions and Rosetta predicted structures, respectively. The last column “*C_α_* RMSD (train)” reports the averaged backbone *C_α_* atom RMSD based on the comparison of each test example to all examples of the same fold and length in the training set, using Rosetta predicted structures.

| Fold | Train | Test | RMSD | $C_\alpha$ RMSD | $C_\alpha$ RMSD (train) |
| --- | --- | --- | --- | --- | --- |
| $\alpha\beta\beta\alpha$ | 92 | 3 | 2.30 | 1.79 | 4.36 |
| $\beta\beta\alpha\beta\beta$ | 533 | 8 | 1.09 | 0.53 | 0.91 |
| $\beta\alpha\beta\beta$ | 787 | 15 | 0.88 | 0.41 | 0.99 |
| $\alpha\alpha\alpha$ | 1330 | 24 | 1.07 | 0.54 | 2.79 |

**Table 2:**
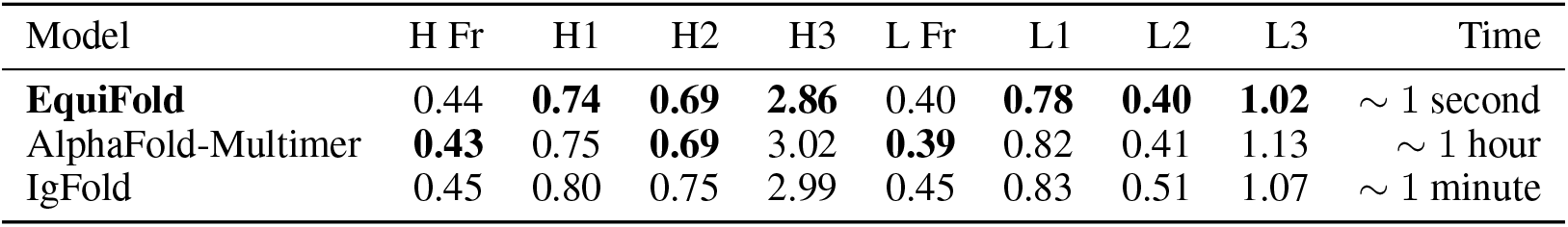
Performance of EquiFold on antibody structures over the test set used in [13] broken down by framework and variable regions. Performance of other models are as reported in [13]. RMSD (in Å) over *N, C_α_, C*, and *O* backbone atoms are reported for valid comparison. Column “Time” reports approximate inference time for EquiFold and approximate run times for other models as reported in [13] for predicting all atom structures.

**Table 3:** The amino acid coarse-graining (CG) scheme used in this work. Each column with “CG” prefix defines a group of atoms belonging to a CG node for each amino acid.

| AA | CG1 | CG2 | CG3 | CG4 |
| --- | --- | --- | --- | --- |
| ALA | $C, C_\alpha, C_\beta, N$ | $C, C_\alpha, O$ | - | - |
| ARG | $C, C_\alpha, C_\beta, N$ | $C, C_\alpha, O$ | $C_\beta, C_\gamma, C_\delta$ | $N_\epsilon, N_{\eta 1}, N_{\eta 2}, C_\zeta$ |
| ASN | $C, C_\alpha, C_\beta, N$ | $C, C_\alpha, O$ | $C_\gamma, N_{\delta 2}, O_{\delta 1}$ | - |
| ASP | $C, C_\alpha, C_\beta, N$ | $C, C_\alpha, O$ | $C_\gamma, O_{\delta 1}, O_{\delta 2}$ | - |
| CYS | $C, C_\alpha, C_\beta, N$ | $C, C_\alpha, O$ | $C_\alpha, C_\beta, S_\gamma$ | - |
| GLN | $C, C_\alpha, C_\beta, N$ | $C, C_\alpha, O$ | $C_\gamma, C_\delta, O_{\epsilon 1}, N_{\epsilon 2}$ | - |
| GLU | $C, C_\alpha, C_\beta, N$ | $C, C_\alpha, O$ | $C_\gamma, C_\delta, O_{\epsilon 1}, O_{\epsilon 2}$ | - |
| GLY | $C, C_\alpha, N$ | $C, C_\alpha, O$ | - | - |
| HIS | $C, C_\alpha, C_\beta, N$ | $C, C_\alpha, O$ | $C_\gamma, C_{\delta 2}, C_{\epsilon 1}, N_{\delta 1}, N_{\epsilon 2}$ | - |
| ILE | $C, C_\alpha, C_\beta, N$ | $C, C_\alpha, O$ | $C_\beta, C_{\gamma 1}, C_{\gamma 2}$ | $C_\beta, C_{\gamma 1}, C_{\delta 1}$ |
| LEU | $C, C_\alpha, C_\beta, N$ | $C, C_\alpha, O$ | $C_\gamma, C_{\delta 1}, C_{\delta 2}$ | - |
| LYS | $C, C_\alpha, C_\beta, N$ | $C, C_\alpha, O$ | $C_\beta, C_\gamma, C_\delta$ | $C_\delta, C_\epsilon, N_\zeta$ |
| MET | $C, C_\alpha, C_\beta, N$ | $C, C_\alpha, O$ | $C_\gamma, C_\epsilon, S_\delta$ | - |
| PHE | $C, C_\alpha, C_\beta, N$ | $C, C_\alpha, O$ | $C_\gamma, C_{\delta 1}, C_{\delta 2}, C_{\epsilon 1}, C_{\epsilon 2}, C_\zeta$ | - |
| PRO | $C, C_\alpha, C_\beta, N$ | $C, C_\alpha, O$ | $C_\beta, C_\gamma, C_\delta$ | - |
| SER | $C, C_\alpha, C_\beta, N$ | $C, C_\alpha, O$ | $C_\alpha, C_\beta, O_\gamma$ | - |
| THR | $C, C_\alpha, C_\beta, N$ | $C, C_\alpha, O$ | $C_\beta, C_{\gamma 2}, C_{\gamma 1}$ | - |
| TRP | $C, C_\alpha, C_\beta, N$ | $C, C_\alpha, O$ | $C_\gamma, C_{\delta 1}, C_{\delta 2}, C_{\epsilon 2}, C_{\epsilon 3}, C_{\zeta 2}, C_{\zeta 3}, C_{\eta 2}, N_{\epsilon 1}$ | - |
| TYR | $C, C_\alpha, C_\beta, N$ | $C, C_\alpha, O$ | $C_\gamma, C_{\delta 1}, C_{\delta 2}, C_{\epsilon 1}, C_{\epsilon 2}, C_\zeta, O_\eta$ | - |
| VAL | $C, C_\alpha, C_\beta, N$ | $C, C_\alpha, O$ | $C_\beta, C_{\gamma 1}, C_{\gamma 2}$ | - |

We define a *reverse* CG mapping 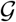 for an amino acid *a_i_* that maps its CG representation to the 3D atom coordinates 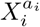 as follows: for each atom in the amino acid, we average the 3D coordinates associated with the atom specified by any of its CG nodes.

### 3.2 Geometric tensor features and initial embedding

To each CG node, we assign a set of geometric tensor features of degree *l* = 0,…, *l_max_* with *n_c_* channels per degree [25, 28, 29]. As there are 2*l* + 1 features associated with an *l*-degree tensor, there are *n_c_* × (*l_max_* + 1)^2^ features in total per node. We define the initial embedding of the nodes 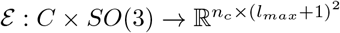 where *C* is the pre-determined set of the CG node types:

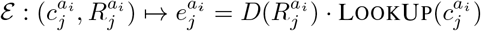

where LookUp: 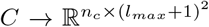 is a typical embedding function and 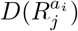 is the direct sum of the Wigner D-matrices *D_l_* corresponding to the tensor features of various degrees *l*:

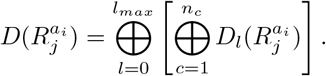

### 3.3 Structure prediction via iterative refinement using an SE(3)-equivariant neural network

Given an input sequence, its CG representation is instantiated with Euclidean transformations whose translations and rotations are sampled from a normal distribution with zero mean and unit variance and a uniform distribution over the SO(3) group, respectively. The output structure is predicted via iterative refinement using an SE(3)-equivariant model that consists of *N_blocks_* blocks sharing the same architecture, where each block is composed of *N_sub_* equivariant sub-blocks. The block architecture is adapted from Equiformer [29], as detailed in Appendix A.2.

Each block takes as input either the initial CG representation or the output of the previous block. A block outputs two *l* = 1 tensors for each node, one of which is used as the vector part of a non-unit quaternion to compute an update *R′* to the node’s rotation *R_in_* and the other to compute an update 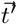 to its translation 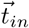 [1] (we drop amino acid and CG node indices for clarity). The input Euclidean transformation 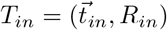 is then updated to 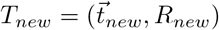 via the following update rules:

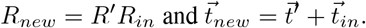

A block can optionally transform or simply copy the input embedding of each node, but either way, it is multiplied by the direct sum of the Wigner D-matrices corresponding to the update rotation *R′*. The proof of equivariance of these update rules is provided in Appendix A.3.

### 3.4 Loss functions

To train the model, the Frame Aligned Point Error (FAPE) loss and the structure violation loss introduced in [1] are computed based on the CG node Euclidean transformations and the reverse CG mapped structures output from each block, respectively. Unlike in [1], all atom FAPE loss is computed for every block. We observed that the structure violation loss is important, since EquiFold is more susceptible to predicting non-physical bond lengths and angles and non-bonded atom distances, due to its modeling of CG nodes in extrinsic 3D space, compared to other models that predict torsion angles representing internal degrees of freedom [1, 2, 3]. However, we do not observe strong instabilities in EquiFold training dynamics like those reported in [1], when the loss is used from the beginning of training. More details on the losses, including their relative weights, other hyper-parameter choices, and training strategies are found in Appendix A.4.

## 4 Results

We report EquiFold’s performance on two structure datasets we curated to focus on structures that present key challenges to protein structure prediction methods: designed mini-proteins and antibody loops. In each challenge, we are primarily predicting structures that are small, have high error in recent blind benchmarks, and have significant potential applications in biotechnology and protein design. These sets also are comprised of regions that elude homology detection and traditional concepts that underpin MSA (as do, by construction, de novo designed sequences). EquiFold achieved high accuracy and speed over these sets, demonstrating that it can enable new downstream protein design and engineering.

### 4.1 *De novo* designed mini-proteins

We first tested EquiFold on a set of de *novo* designed mini-proteins from [10], each having one of four different folds (*ααα, αββα, βαββ*, and *ββαββ*), with associated *in silico* structures predicted using Rosetta [34]. After filtering for sequences with stability score greater than 1, we retained 2,842 sequences whose lengths range from 43 to 50. We randomly split the sequences into train, validation, and test sets of size 2,742, 50, and 50. Test sequences have nearest training sequence similarity ranging from 44% to 80% with mean of 67.2%.^1^ Table 1 shows all atom and *C_α_* RMSD based on the test set broken down by fold. Average inference speed over the entire test set was 0.03 seconds per sequence, with the model containing 1.6M trainable parameters (see Table 4). Figure 2 shows test example predictions overlaid with ground truth structures. Rather than serving as a benchmark relative to other structure prediction models, since the ground truth here are *in silico* predicted structures, this result illustrates the ability of EquiFold to learn a variety of distinct protein topologies with atomic resolution all-atom accuracy.

**Figure 2:**
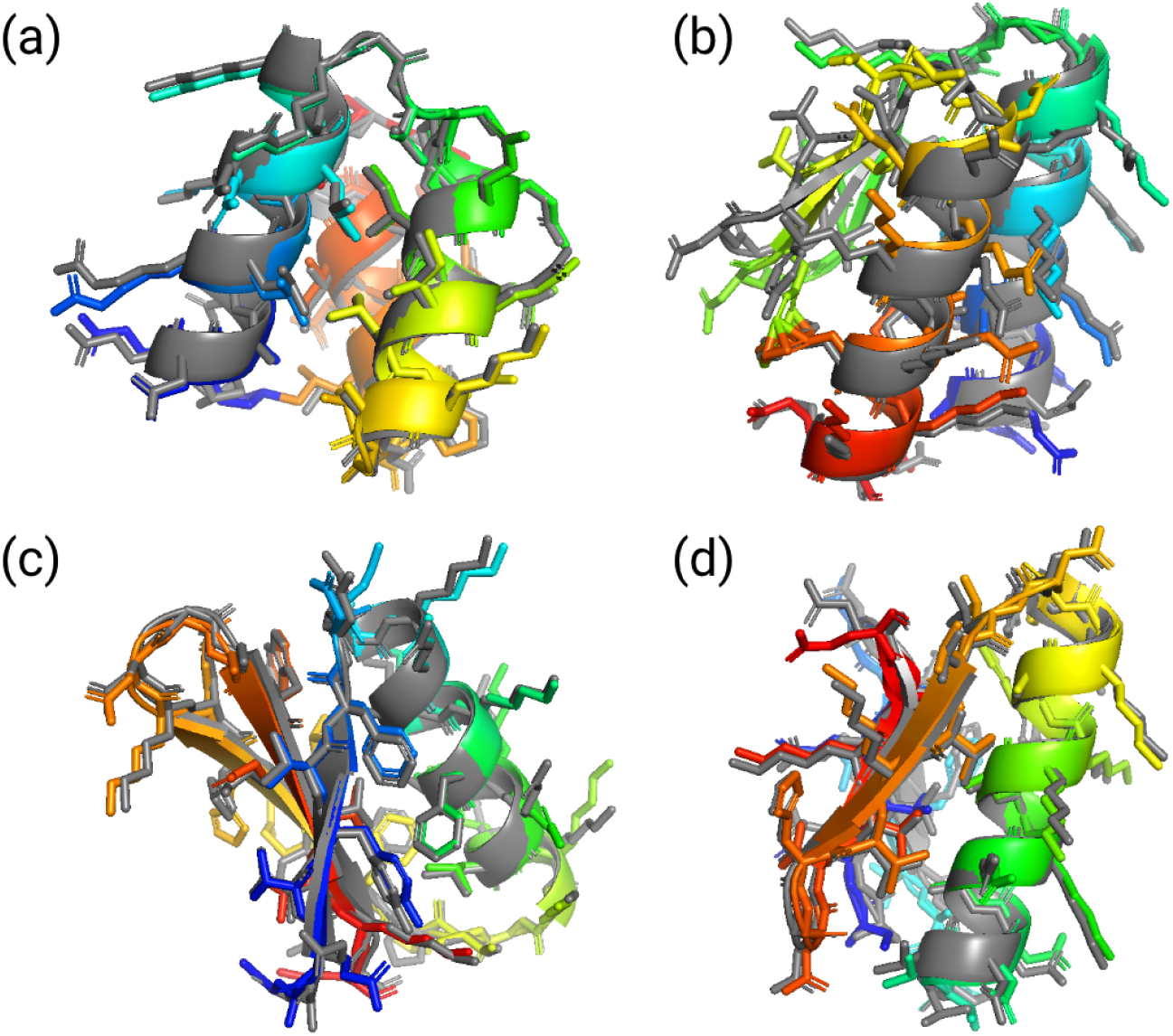
EquiFold’s predictions (rainbow) on test set examples are shown overlaid with the ground truth structures (gray), one for each of the four folds [10]: a) HHH_rd3_0030 (*ααα*, 0.60Å, 62%), b) HEEH_rd2_0948 (*αββα*, 1.50Å, 51%), c) EHEE_rd3_0145 (*βαββ*, 0.86Å, 60%), and d) EE-HEE_rd3_0132 (*ββαββ*, 0.82Å, 60%), where the last two numbers in parentheses correspond to all-atom RMSD and sequence similarity to the nearest training example. EquiFold achieves high accuracy on sequences that have low similarity to any training examples.

**Table 4:** Hyper-parameter values used for the experiments described in Section 4.

| Parameter | Mini-protein | Antibodies |
| --- | --- | --- |
| Number of channels, $n_c$ | 64 | 64 |
| Number of radial basis, $d_{radial}$ | 32 | 32 |
| Number of blocks, $N_{blocks}$ | 4 | 6 |
| Number of sub-blocks, $N_{sub}$ | 3 | 3 |
| Number of attention heads, $N_{head}$ | 2 | 2 |
| Cut-off distance in Å, $r_c$ | 32 | 64 |
| Maximum tensor degree, $l_{max}$ | 1 | 1 |
| Gradient clip value | 1.0 per example | 1.0 per example |
| FAPE loss clip value | 10.0 | 30.0 |
| FAPE loss weight | 1.0 | 1.0 |
| Structure viol. loss weight | 0.5 | 0.2 |
| Number of trainable parameters | 1.6M | 2.4M |

### 4.2 Antibodies

We obtained all antibody (Ab) experimental structures from the PDB [11] that were listed in The Structural Antibody Database (SAbDab) [35], as accessed on January 12, 2022. We processed this dataset to obtain the variable fragment portions of the structures and annotated the sequences with Chothia numbering using ANARCI [36]. We obtained 6,789 structures with a resolution better than 4Å and deposited before July 1, 2021 as training set, of which 50 structures were used for validation. We used the same test set structures as in [13] that have resolution better than 3 Å and deposited after the aforementioned date. Compared to other models, EquiFold achieves similar or better accuracy in backbone atom RMSD across different sequence regions (see Table 2) and results in all atom RMSD of 1.33 Å averaged over the test set. Importantly, the model has fast inference speeds of approximately 1 second per Ab on average on a single A100 GPU for predicting all atom structures and contains 2.4M trainable parameters (see Table 4), compared to 27.6M of IgFold (including 26M of the antibody language model AntiBERTy) [13] and 93.2M of AlphaFold [1] that requires time-consuming input preparation steps. Given its high speed and accuracy, it is practical to predict structures at high-throughput scale for millions of antibodies observed in deep sequencing data sets [37] and integrate the model in a design workflow [16, 21]. Fig 3 shows selected test set predictions overlaid with ground truth structures.

**Figure 3:**
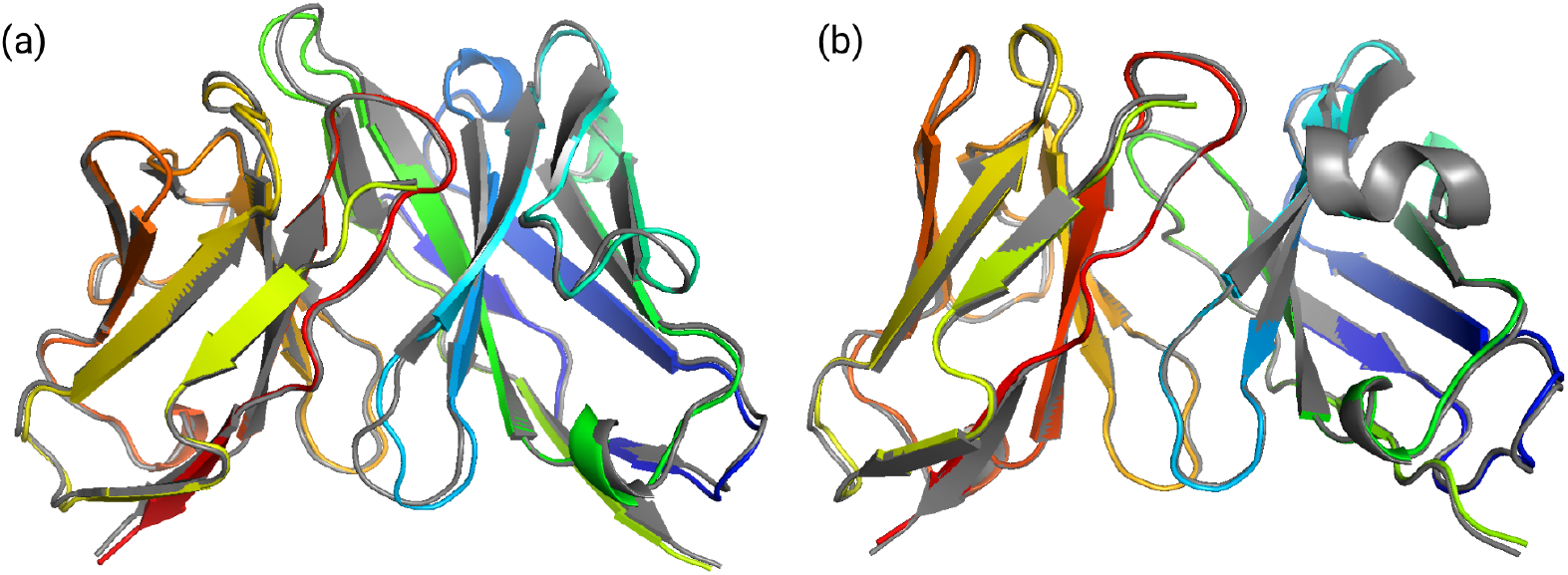
EquiFold’s predictions (rainbow) on two examples from the test set [13] are shown overlaid with the ground truth structures (gray): a) 71fb_HL (1.09Å, 71%) and 7oo2_CD (0.98Å, 72%), where the two numbers in parentheses correspond to all-atom RMSD and sequence similarity to the nearest training example.

## 5 Conclusion and Future Work

We introduced EquiFold, a new end-to-end differentiable protein structure prediction model that uses a novel coarse-grained (CG) representation of proteins. The model achieves high test set accuracy on two datasets, while running at a substantially faster speed. Notably, it is trained on significantly less data and does not rely on multiple sequence alignments [1, 2] or protein language model embeddings [3, 4, 13]. Its accuracy and speed make it practical to integrate EquiFold as a sub-component of a larger model. For instance, EquiFold can be combined in an end-to-end differentiable fashion with another neural network that predicts various molecular properties based on the structure output by its last block.

We leave to future work training EquiFold on more general classes of proteins from the PDB [11] and examining its generalizability to novel folds unseen in training data. Scaling to larger proteins will require addressing the quadratic complexity of the message passing layers, possibly using similar strategies in earlier works [1]. To integrate physical priors, we will extend the CG representation to include hydrogens and implement various energy functions such as the Rosetta all-atom energy function [34]. With such utilities, EquiFold can be adapted to generate conformational ensembles within an energy band, perform flexible docking, and fit structural models to experimental data such as that from cryogenic electron microscopy (cryo-EM) and hydrogen-deuterium exchange (HDX). Lastly, the CG representation could be used in generative modeling of protein sequence and structure together, rather than modeling them sequentially as done in recent works [16, 20, 21].

## A Appendix

### A.1 Computing coarse-grained node template coordinates and ground-truth Euclidean transformations

For each coarse-grained (CG) node defined in Table 3, we compute the template coordinates of all the atoms comprising the node as follows. Given a protein structure dataset 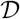, we compute the Euclidean transformation *T_q_* corresponding to each CG node *q* in 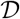 using Algorithm 21 of [1] with the 3D coordinates of the first three atoms of the CG node *q*’s atom group as input. Next, we apply the inverse Euclidean transformation 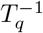 to the 3D coordinates of all the atoms in the node q into the corresponding local frame. Lastly, we average the observed coordinates in the local frames across all instances in the dataset 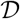 grouped by CG node type defined in Table 3 and subtract their geometric centroids.

To compute the ground truth Euclidean transformation for a given CG node to be used in the FAPE loss [1], we use the Kabsch algorithm [38] to determine the transformation from the template coordinates to the observed coordinates that minimizes the root mean square error (RMSE) of the node’s constituent atoms. To elaborate, for the *j*-th CG node 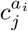 containing *M* atoms of the i-th amino acid *a_i_*, we apply the Kabsch algorithm to the template coordinates 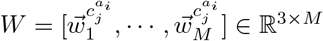 and the corresponding input coordinates 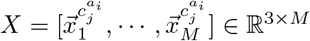. The algorithm factorizes the covariance matrix 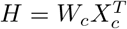 into eigenvectors and values using singular vector decomposition,

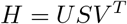

where *W_c_* and *X_c_* are mean centered template and input coordinates. The resulting rotation and translation are

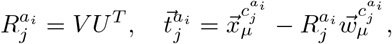

where 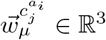 and 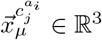 are the mean coordinates of *W* and *X* respectively.

### A.2 SE(3)-equivariant neural network architecture details

Each block of the neural network consists of *N_sub_* sub-blocks that share the same architecture, where each sub-block is an adapted version of the Equiformer’s “Transformer block” [see 29, Figure 1]. As the general theory of SE(3)-equivariant neural networks and the implementation of the Transformer block are well-described in [29], here we describe only the modifications in our adapted version.

Given the input set of coarse-grained (CG) nodes and their Euclidean transformations, the block initially computes pairwise distances *r_ij_* and normalized distance vectors 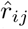, where *i* and *j* index CG nodes. *r_ij_* is projected onto *d_radial_* sinusoidal radial bases with learnable weights and a cutoff distance *r_c_*, which are used in radial functions that parameterize tensor products in the Equivariant Graph Attention module. 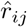 is used to compute spherical harmonics 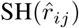 input to tensor products. When training the network, gradients do not propagate through *r_ij_* and 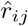.

In our adapted version of the Equivariant Graph Attention module, after the application of the initial linear layers to input node embeddings, instead of element-wise summation, channel-wise fully connected tensor products are applied to the embeddings of every CG-node pair *ij*, which is followed by another linear layer to produce output tensors *x_ij_* with the same number of channels as the input. Next, Depth-wise Tensor Product (DTP) is applied to *x_ij_* and 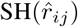 with a radial function that takes as input the radial bases mentioned above and additionally a scalar *edge* embedding vector corresponding to the primary amino acid sequence distance for the CG-pair *ij*, clamped at maximum absolute distance of 32; for input proteins with multiple chains, sequence distances across chains are set at the maximum distance. The edge embedding is implemented via a simple look-up table with learnable weights and has the same dimension as the number of channels in input tensors. The output of the DTP layer is uniformly shuffled and grouped by *N_head_* attention heads and a linear layer is applied to produce tensors of various degrees with appropriate channel numbers for the remainder of the module.

The output of each sub-block except for the last one are updated node embeddings corresponding to the input CG nodes. The last sub-block outputs only two *l* = 1 vectors per CG node as mentioned in Section 3.3. Edge embeddings are shared across sub-blocks of a given block.

### A.3 Proof of equivariance of the Euclidean transformation update

Under a global Euclidean transformation 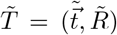, the Euclidean transformation 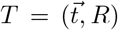 corresponding to a CG node that specifies the mapping between the observed and template coordinates transforms as

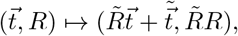

and the update transformation 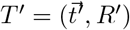 output by a block transforms as

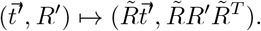

We show that the update rules given in Section 3.3 are equivariant by applying a global transformation 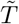 to the input 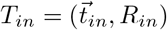 and update *T*′ transformations and showing that their composition is equivalent to the global transformation of the output 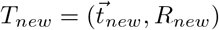:

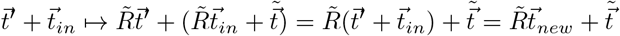

and

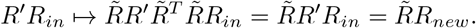

### A.4 Hyper-parameters and training details

Table 4 provides the hyper-parameter values used for the two experiments described in Section 4. Both models were trained using the Adam optimizer [39] using PyTorch [40] with `beta=(0.9, 0.999)` and `weight_decay=`10^-8^. We used a warm-up phase of 1,000 steps, where the learning rate was linearly increased from 0 to 10^-3^ and input structures to the model were linearly interpolated between the ground truth structures and random initializations; for rotations, we used quaternion spherical linear interpolation [20]. After the warm-up phase, the learning rate was decreased to 10^-4^ following a cosine annealing schedule over 100,000 steps and was kept at the final value thereafter.

Both models were trained with the FAPE and structure violation losses computed on the output structure of each block. The structure violation loss was modified to penalize deviations from standard values by more than 3 standard deviations, compared to the 12 standard deviations used in [1]. A mini-batch size of 8 was used with 8 A100 GPUs in PyTorch’s Distributed Data Parallel mode [40]. Model training for the miniprotein experiment was stopped after one day as the validation loss converged. The antibody model was trained for approximately 20 days.

To circumvent large memory requirement originating from the quadratic computational complexity of attention layers, we computed gradient updates of the learnable weights of the models on a block-by-block basis, stopping gradient propagation through rotations and translations after each block. Early experiments on mini-proteins showed that this model weight update algorithm did not substantially affect training dynamics. In addition, we implemented a distinct embedding layer for each block. Similarly, we did not observe a significant benefit to using tensors of degrees higher than 1. We leave to future work a more careful benchmark of different model weight update schemes and other hyper-parameters.

## Footnotes

1 Sequence similarity is defined as (*l_query_ – n_edit_*)/*l_query_* where *l_query_* is the length of the query sequence and *n_edit_* the Levenshtein edit distance between query and target sequences.

## References

[1] J. Jumper et al. Highly accurate protein structure prediction with AlphaFold. Nature, 2021.

[2] M. Baek et al. Accurate prediction of protein structures and interactions using a three-track neural network. Science, 2021.

[3] R. Wu et al. High-resolution de novo structure prediction from primary sequence. bioRxiv, 2022.

[4] Z. Lin et al. Language models of protein sequences at the scale of evolution enable accurate structure prediction. bioRxiv, 2022.

[5] R. Chowdhury et al. Single-sequence protein structure prediction using language models from deep learning. bioRxiv, 2021.

[6] A. Meller et al. Predicting the locations of cryptic pockets from single protein structures using the PocketMiner graph neural network. bioRxiv, 2022.

[7] P. Gainza et al. Deciphering interaction fingerprints from protein molecular surfaces using geometric deep learning. Nat Methods 2020.

[8] M. L. Fernández-Quintero et al. Paratope states in solution improve structure prediction and docking. Structure, 2022.

[9] B. Jing, S. Eismann, P. N. Soni, and R. O. Dror. Equivariant graph neural networks for 3d macromolecular structure. arXiv, 2022.

[10] G. J. Rocklin et al. Global analysis of protein folding using massively parallel design, synthesis, and testing. Science, 2017.

[11] H. Berman, K. Henrick, and H. Nakamura. Announcing the worldwide Protein Data Bank. Nat Struct Mol Biol, 2003.

[12] J. A. Ruffolo, J. Sulam, and J. J. Gray. Antibody structure prediction using interpretable deep learning. Patterns, 2022.

[13] J. A. Ruffolo, L.-S. Chu, S. P. Mahajan, and J. J. Gray. Fast, accurate antibody structure prediction from deep learning on massive set of natural antibodies. bioRxiv, 2022.

[14] R. Das. Four small puzzles that rosetta doesn’t solve. PLOS ONE, 2011.

[15] C. Hsu et al. Learning inverse folding from millions of predicted structures. bioRxiv, 2022.

[16] J. Dauparas et al. Robust deep learning based protein sequence design using ProteinMPNN. Science, 2022.

[17] K. T. Simons et al. Ab initio protein structure prediction of CASP III targets using ROSETTA. Protein, 1999.

[18] A. Del Vecchio et al. Neural message passing for joint paratope-epitope prediction. arXiv, 2021.

[19] V. Gligorijević et al. Structure-based protein function prediction using graph convolutional networks. Nat Commun. 2021.

[20] N. Anand and T. Achim. Protein structure and sequence generation with equivariant denoising diffusion probabilistic models. arXiv, 2022.

[21] J. Wang et al. Scaffolding protein functional sites using deep learning. Science, 2022.

[22] S. Batzner et al. E(3)-equivariant graph neural networks for data-efficient and accurate interatomic potentials. Nat Commun, 2022.

[23] I. Batatia et al. The Design Space of E(3)-Equivariant Atom-Centered Interatomic Potentials. arXiv, 2022.

[24] I. Batatia et al. MACE: Higher order equivariant message passing neural networks for fast and accurate force fields. arXiv, 2022.

[25] N. Thomas et al. Tensor field networks: Rotation- and translation-equivariant neural networks for 3D point clouds. arXiv, 2022.

[26] F. B. Fuchs et al. SE(3)-Transformers: 3D Roto-Translation Equivariant Attention Networks. arXiv, 2020.

[27] V. G. Satorras, E. Hoogeboom, and M. Welling. E(n) equivariant graph neural networks. arXiv, 2022.

[28] J. Brandstetter et al. Geometric and physical quantities improve E(3) equivariant message passing. arXiv, 2022.

[29] Y.-L. Liao and T. Smidt. Equiformer: Equivariant graph attention transformer for 3d atomistic graphs. arXiv, 2022.

[30] O.-E. Ganea et al. Independent SE(3)-equivariant models for end-to-end rigid protein docking. In International Conference on Learning Representations, 2022.

[31] W. Jin, D. Barzilay, and T. Jaakkola. Antibody-antigen docking and design via hierarchical structure refinement. In Proceedings of the 39th International Conference on Machine Learning, 2022.

[32] N. Frey et al. Neural Scaling of Deep Chemical Models. chemRxiv, 2022.

[33] A. Musaelian et al. Learning local equivariant representations for large-scale atomistic dynamics. arXiv, 2022.

[34] J. K. Leman, et al. Macromolecular modeling and design in Rosetta: recent methods and frameworks. Nat Methods, 2020.

[35] C. Schneider, M. Raybould, and C. M. Deane, SAbDab in the age of biotherapeutics: updates including SAbDab-nano, the nanobody structure tracker, Nucleic Acids Research, 2022.

[36] J. Dunbar and C. M. Deane. ANARCI: antigen receptor numbering and receptor classification. Bioinformatics, 2016.

[37] T. H. Olsen, F. Boyles, and C. M. Deane. Protein Science, 2021.

[38] W. Kabsch. A solution for the best rotation to relate two sets of vectors. Acta Crystallographica, 1976.

[39] D. P. Kingma and J. Ba. Adam: A method for stochastic optimization. arXiv, 2017.

[40] A. Paszke et al. PyTorch: An imperative style, high-performance deep learning library. arXiv, 2019.

